# G protein-Coupled Receptor Associated Sorting Protein GASP1 mediated trafficking of the Glucagon Like Peptide-1 Receptor contributes to the development of tolerance to incretin drugs

**DOI:** 10.1101/2024.08.09.607397

**Authors:** Anirudh Gaur, Caroline M Keeshen, Madeline King, Noah Dirks, Marcus Flisher, Bethany P Cummings, Mark O Huising, Jennifer L Whistler

## Abstract

Incretin mimetic drugs are in widespread use for the treatment of type 2 diabetes and obesity and more recently have been prescribed for weight loss in otherwise healthy individuals. These drugs are all agonists of the glucagon-like peptide 1 receptor (GLP-1R) and function by supplementing effects produced by the endogenous hormone agonist glucagon-like peptide 1 (GLP-1). The therapeutic benefits of these medications, including improved glucose control and weight loss, require continued usage and wane with time. The molecular mechanisms underlying this loss of effect to incretin drugs remain unknown. Following activation by agonist and signaling to G protein, the GLP-1R engages arrestins and is endocytosed. Here we investigated the role of G protein-coupled receptor associated sorting protein 1 (GASP1), a critical regulator of the post-endocytic trafficking of GLP-1R, on tolerance to GLP-1R agonist drug. We found that tolerance to incretin drug was prevented at the cellular, tissue and whole animal level in mice with a selective disruption of the GASP1 protein in beta cells of the pancreatic islet. These studies implicate post-endocytic sorting of the GLP-1R in the loss of effectiveness of incretin therapeutics with prolonged use. These findings also suggest a novel strategy to prevent tolerance by biasing incretin drugs for G protein and away from arrestin engagement.

## Introduction

Type 2 diabetes and obesity present significant global public health challenges, with rising prevalence and associated complications^1–3^. In recent years, incretin drugs have emerged as effective therapeutics for managing both conditions^4–7^. These medications include memetics of the hormone glucagon-like peptide-1 (GLP-1) which activate the GLP-1 receptor (GLP-1R) as well as inhibitors of dipeptidyl peptidase-4 (DPP-4) which breaks down endogenous GLP-1^8–10^. The efficacy of these drugs reflects their ability to harness the body’s own incretin system to enhance insulin secretion and regulate blood glucose levels, even in subjects where these processes have been diminished^11,12^. Clinical trials have consistently demonstrated that GLP-1R agonist drugs including Byetta^®^ (Exenatide/exendin-4), Victoza^®^ (liraglutide) and Ozempic^®^ (semaglutide) produce significant reductions in HbA1c levels and fluctuations in postprandial glucose excursions^13–18^. Additionally, these drugs promote weight loss, as they delay gastric emptying, promote satiety and have central effects on appetite regulation^19–21^.

One notable challenge that may arise with long-term use of incretin drugs is the development of “tolerance” defined as reduced therapeutic response to incretin drugs over time^22–25^. Elucidating the mechanisms responsible for tolerance could inform new strategies to mitigate its impact and optimize the long-term effectiveness of incretin drugs in type 2 diabetes and obesity management. In addition, tolerance to long-term use of these therapeutics may also have significant new implications, as these drugs are now in widespread use for weight loss in otherwise healthy individuals^26^.

The GLP-1R is a class B G protein-coupled receptor (GPCR) expressed in pancreatic islet cells, the gastrointestinal tract and the central nervous system^27,28^. GLP-1 and GLP-1 memetics initiate a cascade of intracellular signaling events via G protein signaling when they are bound to the GLP-1R. In beta cells, this G protein signaling enhances glucose-stimulated insulin secretion and is known as the “Incretin effect”^29,30^. Following G protein activation, signaling from most GPCRs including the GLP-1R is attenuated by desensitization events that includes phosphorylation by GPCR kinases (GRKs) and recruitment of arrestins, which arrest G protein signal and promote GPCR endocytosis^30–33^. Endocytosed receptors are then sorted into one of two main pathways, the recycling pathway whereby they are returned to the plasma membrane or the degradative pathway whereby they are sent to the lysosome for degradation^34^. The post-endocytic fate of a GPCR has profound implications for signal transduction, especially under conditions, such as exogenous drug use, where there are high concentrations of ligand and therefore a large degree of receptor endocytosis^35^. For receptors that are recycled, endocytosis serves to rapidly re-sensitize signal transduction while for receptors that are degraded, endocytosis will promote prolonged loss of signaling^36^.

The GLP-1R has been shown to undergo both recycling and degradation after endocytosis depending on the cell type^32,37^. The GLP-1R has also been shown to interact with GPCR-associated sorting protein 1 (GASP1, encoded by the *GPRASP1* gene), a critical mediator of the post-endocytic targeting of numerous GPCRs to the lysosome^38–40^. We hypothesized that tolerance to GLP-1R agonists could be mediated by interaction with GASP1 and consequent post-endocytic targeting of the GLP-1R to the lysosome. Specifically, we hypothesized that with high and chronic agonist exposure from the memetic drugs, GLP-1R synthesis would be insufficient to keep up with post-endocytic degradation of the GLP-1R and effectiveness of the drug would diminish. Here we examine this hypothesis in INS-1 cells, pancreatic islets, and mice *in vivo*, with and without functional GASP1 protein. We found that deletion of GASP1 prevents tolerance to chronic incretin drug in all three systems. These findings implicate post-endocytic trafficking of the GLP-1R as a mechanism that limits the efficacy of incretin therapeutics during chronic use. These studies also suggest that an incretin memetic drug that promotes G protein signaling but not arrestin engagement could provide prolonged therapeutic utility.

## Results

### Pre-treatment of INS-1 cells with exendin-4 decreases GLP-1R signaling in GASP1-WT but not in GASP1-KO cells

To examine the role of GASP1 in modulating the activity of GLP-1R in response to chronic incretin drug, we utilized CRISPR-Cas9 to delete the GASP1 gene (*Gprasp1*) in INS-1 cells (GASP1-KO), which was confirmed by immunoblot (Fig. 1A). GASP1-WT and GASP1-KO cells were then pretreated for 3-hours with either exendin-4 (Ex-4) or vehicle and GLP-1R function was assessed both at the level of cAMP accumulation and insulin secretion (see schematic Fig. 1B). GASP1-WT and GASP1-KO cells pretreated with vehicle showed equivalent dose-dependent increases in cAMP levels, indicating that deletion of GASP1 had no effect on GLP-1R function in response to acute Ex-4 (Fig. 1C vs. 1D, E_max_ GASP1-WT: 100%, GASP1-KO: 95.14% ± 3.26, EC_50_: 0.59 ± 0.28 and 0.55 ± 0.26 nM respectively Table S1). GASP1-WT cells pretreated with Ex-4 showed a significantly reduced cAMP response to Ex-4 compared to vehicle treated GASP1-WT cells (Fig. 1C, E_max_: 100% vs. 44.30% ± 11.53, **p = 0.008, Table S1) indicating that these cells developed tolerance to Ex-4. In contrast, GASP1-KO cells pretreated with Ex-4 showed comparable cAMP responses to vehicle pretreated cells (Fig. 1D, E_max_: 95.14% ± 3.26 vs. 92.17% ± 4.32, Table S1). GASP1-WT and GASP1-KO cells pretreated with vehicle also showed equivalent dose-dependent insulin secretion in response to Ex-4 (Fig. 1E and F, E_max_: 100% & 99.98% ± 4.76, EC_50_: 0.044 ± 0.01 & 0.035 ± 0.02 nM respectively, Table S2).

**Figure 1:**
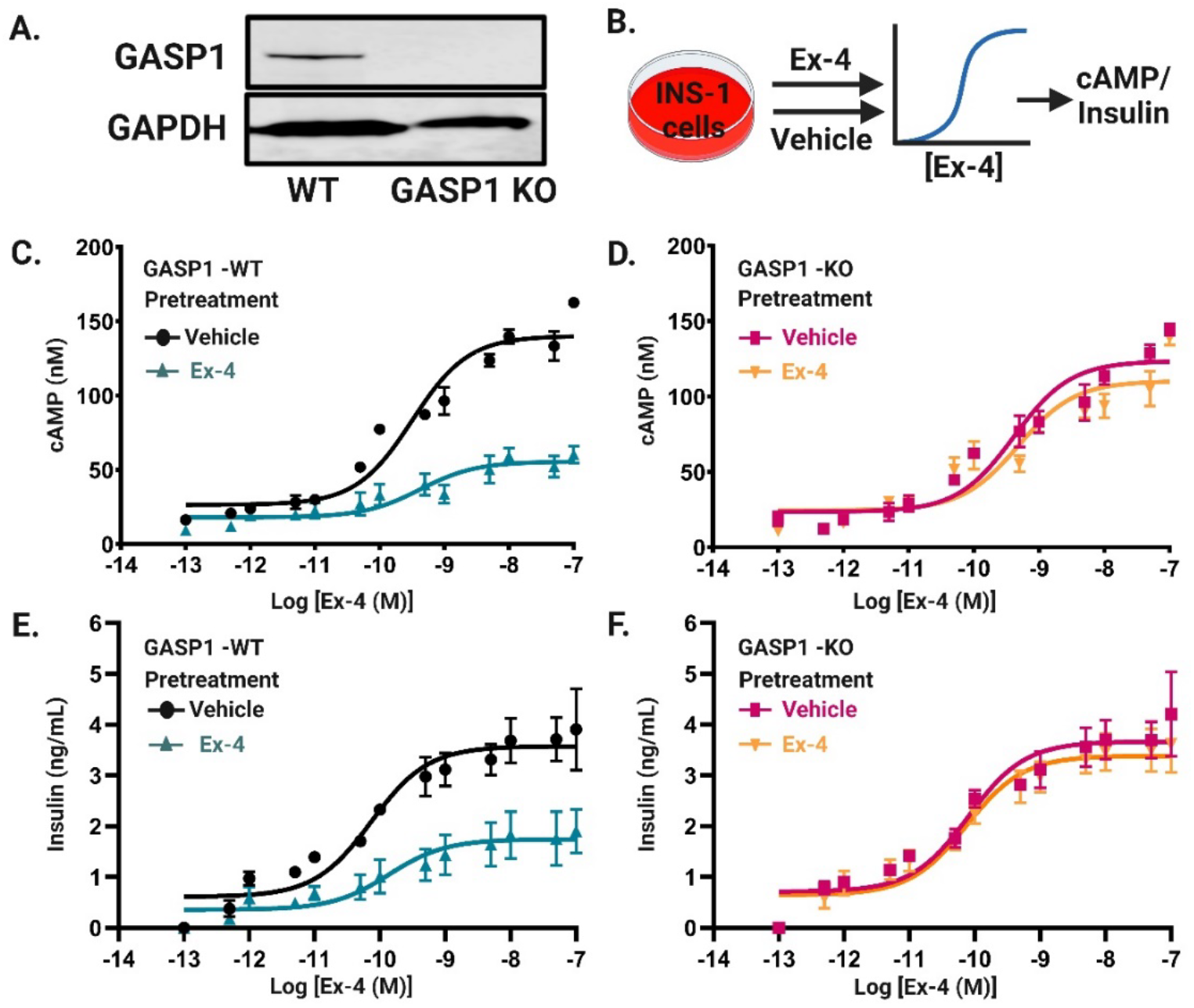
Pre-treatment of INS-1 cells with Ex-4 decrease GLP-1R signaling in GASP1-WT but not in GASP1-KO cells. A) Immunoblot detection of GASP1 protein (upper blot) and GAPDH protein (lower blot) in wild-type (WT) and GASP1 knockout (GASP1-KO) INS-1 cells. B) Schematic of experimental approach for cAMP and insulin release assays. Briefly, both WT and GASP1-KO INS-1 cells were pretreated with either exendin-4 (Ex-4,100nM) or vehicle for 3-hours, washed and then a dose response for cAMP and insulin release in response to Ex-4 was measured. C, D) Dose-response curves showing Ex-4-induced changes in cAMP accumulation in WT INS-1 cells (C) and GASP1-KO INS-1 cells (D) with or without Ex-4 pretreatment. The curves are representative of 3 different experiments performed on different days with each dose in triplicate. Error bars represent S.E.M. Average E_max_ and EC_50_ from all three experiments are shown in Supp. Table S1. WT but not GASP1-KO cells show reduced response to Ex-4 (E_max_ for vehicle vs Ex-4 pretreated for GASP1-WT cells: 100% and 44.30%±11.53, *p = 0.008; for GASP1-KO cells: 95.14±3.26 and 92.17±4.32, ns). E, F) Dose-response curves showing Ex-4 dependent insulin release by WT INS-1 cells (E) and GASP1-KO INS-1 cells (F) with or without Ex-4 pretreatment. Shown is an average of 3 different experiments performed on different days in triplicate. Error bars represent S.E.M. E_max_ and EC_50_ are shown in Supp. Table S2. WT but not GASP1-KO cells show reduced response to Ex-4 (E_max_ for vehicle vs Ex-4 pretreated for GASP1-WT cells: 100% and 56.65±13.14, *p = 0.03; for GASP1-KO cells: 99.98%±4.76 and 92.49±3.34, ns).

Following Ex-4 pretreatment, GASP1-WT but not GASP1-KO cells showed tolerance to Ex-4-dependent insulin secretion compared with vehicle pretreated cells (Figure 1E and F; E_max_: 100% vs. 56.65% ± 13.14 for WT, *p = 0.03, E_max_: 99.98% ± 0.02 vs. 92.49 ± 0.02 for KO, Table S2). Together these data demonstrate that GASP1 is necessary for the development of tolerance to Ex-4 in INS-1 cells.

### Chronic Ex-4 abolishes GLP-1R-mediated insulin secretion from WT but not β-GASP1-KO islets

We next used *ex vivo* mouse pancreatic islets to determine if they also exhibit GASP1-dependent tolerance to Ex-4. We generated mice with a floxed-GASP1 allele (fl-GASP1, Supp. Fig. 1)^41^. We then crossed the fl-GASP1 mice to mice with Cre-recombinase driven by the beta-cell specific urocortin-3 gene promoter (UCN3-Cre)^42^, producing mice with islet β-cell-specific knock out of GASP1 (β-GASP1-KO) (Fig. 2A). The deletion of GASP1 was confirmed with qPCR analysis of islets isolated from GASP1-WT and β-GASP1-KO mice, which revealed a significant reduction in GASP1 mRNA expression in β-GASP1-KO islets (**p<0.01) compared to GASP1-WT islets (Fig. 2B). Deletion of GASP1 also led to a significant reduction in GLP-1R gene expression in β-GASP1-KO islets (*p<0.05) compared to GASP1-WT islets, possibly because GLP-1R synthesis needs were reduced due to prevention of post-endocytic GLP-1R degradation.

**Figure 2:**
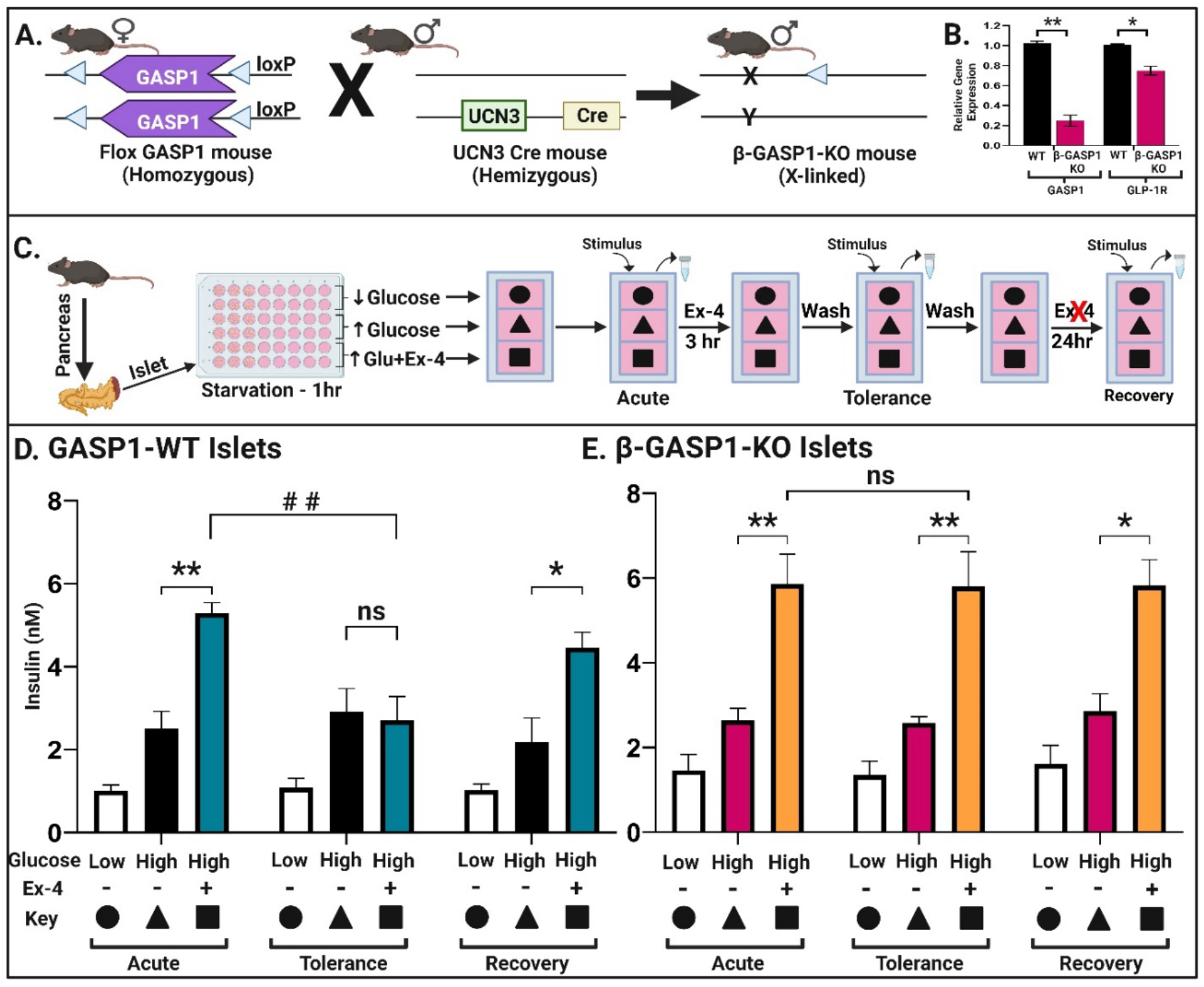
Chronic Ex-4 abolishes GLP-1R mediated insulin secretion in WT but not β-GASP1-KO islets. A) Schematic of strategy used to generate β-cell specific GASP1 knockout mice (β-GASP1-KO). β-GASP1-KO mice were created by crossing female homozygous floxed-GASP1 mice with male heterozygous UCN3-Cre mice. Male littermates with and without the UCN3-Cre driver were used for all experiments. B) qPCR analysis of GASP1 and GLP-1R expression in WT and β-GASP1-KO islets. The bars represent the mean data from 2 biological replicates, each with 3 technical replicates. Data were normalized to the endogenous housekeeping gene hypoxanthine phosphoribosyltransferase 1 and are expressed as relative gene expression compared to WT islets. *p<0.05, **p<0.01, WT vs. β-GASP1-KO, unpaired t-test. C) Schematic of the longitudinal islet insulin secretion assay. The islets were monitored for insulin secretion acutely, following 3-hours of Ex-4 pretreatment and again after 24-hours of recovery. D, E) Insulin secretion from islets of WT (D) and β-GASP1-KO (E) islets in response to low glucose (5.5mM), high glucose (11mM) and high glucose + Ex-4 acutely (Left panels), after 3-hours of Ex-4 treatment (middle panels) and after 24-hours of recovery in regular media (right panels). The bars show mean data of three biological replicates, each comprised of three technical replicates. Error bars represent S.E.M. WT, but not β-GASP1-KO islets show reduced Ex-4 response after Ex-4 pretreatment. *p<0.05, **p<0.01, ***p<0.001, high glucose vs. high glucose + Ex-4, and ^##^p<0.01 Acute vs. Ex-4 pretreatment determined by two-way ANOVA.

We next utilized a longitudinal insulin secretion paradigm on islets isolated from GASP1-WT and β-GASP1-KO mice to examine tolerance to Ex-4 (Fig. 2C). Insulin secretion from these islets was assayed at three timepoints (Fig. 2C, Acute, Tolerance, Recovery). For the first timepoint, islets were treated with low glucose (5.5mM), high glucose (11mM) and high glucose+Ex-4 (100nm) for 1-hour, and then a sample of media was removed and assayed for insulin. Both GASP1-WT and β-GASP1-KO islets displayed robust insulin secretion when stimulated with high glucose (Fig. 2D and E, white vs. black and white vs. magenta bar respectively, “Acute”). Stimulation of both GASP1-WT and β-GASP1-KO islets with high glucose and 100nM Ex-4 produced significant increases in insulin secretion compared to high glucose alone (Fig. 2D and E, black vs. teal and magenta vs. orange bar respectively, “Acute”) demonstrating an equivalent acute incretin effect in GASP1-WT vs. β-GASP1-KO islets. Thus, neither deletion of GASP1 nor reduction in GLP-1R synthesis in the β-GASP1-KO islets affected acute responses to Ex-4.

We next treated these islets with Ex-4 for 3-hours, followed by a wash and 15 minutes recovery. Islets were then incubated with low glucose, high glucose, or high glucose + Ex-4 for 1-hour and media once again assayed for insulin secretion. GASP1-WT islets showed a robust response to high glucose indistinguishable from the acute response, however, they showed no response to Ex-4 (Fig. 2D, middle panel, black vs teal bar, “Tolerance”). In contrast, β-GASP1-KO islets maintained their incretin effect following the 3-hours Ex-4 pretreatment (Fig. 2E, middle panel, magenta vs orange bar). These results suggest that WT-GASP1, but not β-GASP1-KO islets, develop tolerance to the incretin effect under constant incretin exposure. Importantly, this effect was not permanent, as both GASP1-WT and β-GASP1-KO islets showed a substantial incretin effect after 24-hours of recovery in the absence of Ex-4 (Fig. 2D and E, right panel, “Recovery”). These data indicate that GASP1 plays a crucial role in modulating GLP-1R function and the development of tolerance to Ex-4 in mouse islets.

### Wild-type mice develop tolerance to exendin-4

We next investigated the response to chronic Ex-4 treatment *in vivo*. We implemented a 9-week longitudinal paradigm and assayed both glucose clearance following oral gavage and glucose-stimulated insulin secretion (GSIS) at multiple timepoints in WT mice treated with chronic saline or Ex-4 (see Fig. 3A for a schematic of experimental design; see supp Fig. 2 for schematic of blood collection timepoints). At baseline (week 1), glucose produced robust insulin secretion (Fig. 3B, solid vs hatched bar; **p<0.01 baseline vs glucose) and was cleared within 120 minutes (Supp. Fig. 3B). At week 2, WT mice treated with Ex-4 (200 μg/kg) showed enhanced insulin secretion compared to the saline-treated group (Fig. 3C solid black vs. teal bar; ^##^p<0.01 glucose + saline vs. glucose + Ex-4), demonstrating the incretin effect. Mice were then treated twice daily with either saline or Ex-4 for an additional 7 weeks. At week 3, we evaluated GSIS (Fig. 3D) and glucose clearance (Supp. Fig. 3D) and found that response to Ex-4 was indistinguishable from week 2. In contrast, at week 8 (6 weeks of Ex-4), we found that GSIS in response to glucose + Ex-4 was indistinguishable from that produced by glucose + saline (Fig. 3E, solid black vs. teal bar, ns), indicating the development of tolerance to Ex-4. Lastly, at week 9, we examined GSIS in response to glucose alone to determine the response to endogenous incretins (Fig. 3F). These data show that mice treated chronically with Ex-4 show both a reduction in their response to exogenous incretin (Fig. 3G, week 3 vs. week 8) and to oral glucose alone which includes the endogenous incretin effect (Fig. 3G, week 1 vs. week 9).

**Figure 3:**
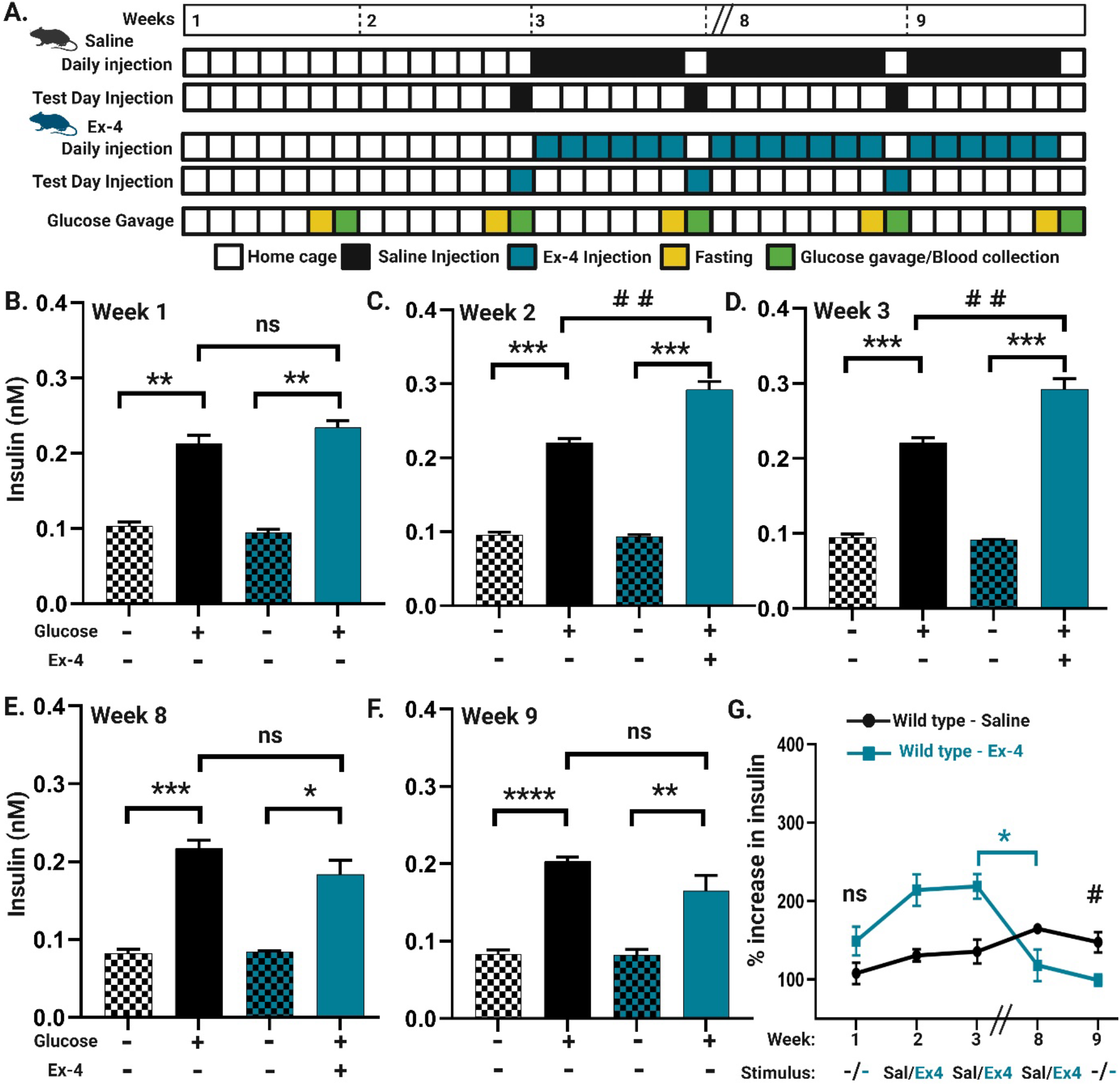
Wild-type mice develop tolerance to incretin. A) Schematic of the longitudinal mouse paradigm for plasma insulin measurement in WT mice repeatedly injected with saline or Ex-4 (200μg/Kg). B) Glucose stimulated insulin secretion (GSIS) in WT mice (n=8) following glucose gavage at baseline (Week 1). Mice were administered glucose orally (2g/kg body weight) and plasma insulin was measured. Mice show robust GSIS in response to glucose (hatched vs non-hatched bar, **p<0.01). C) GSIS in WT mice following acute Ex-4 treatment (Week 2). Mice received either Ex-4 (n=4) or saline (n=4) injection on test day, followed by oral glucose administration and plasma insulin measurement. WT mice treated with Ex-4 show enhanced insulin secretion compared to saline treated mice (black vs teal bar, ^##^p<0.01). D) GSIS in WT mice following one week of twice daily Ex-4 or saline (Week 3). Mice treated with Ex-4 maintain an Ex-4-dependent incretin effect compared to saline treated mice (black vs. teal bar, ^##^p<0.01). E) GSIS in WT mice after six weeks of chronic daily Ex-4 or saline treatment (Week 8). Mice continue to receive twice daily injections of either Ex-4 or saline for six weeks. Mice treated with Ex-4 show loss of the Ex-4-dependent incretin effect (Tolerance) resulting in comparable insulin production as that of saline treated mice (black vs. teal bar). F) GSIS in WT mice after seven weeks of chronic Ex-4 or saline treatment (Week 9). Mice continued to receive twice daily injections of either Ex-4 or saline. On test day, plasma insulin concentration was measured after oral glucose administration without either Ex-4 or saline injections. GSIS in Ex-4 treated mice is significantly lower than saline treated mice (black vs. teal bar). The bar graphs (B-F) represent the mean ± S.E.M from multiple mice. *p<0.05, **p<0.01, ***p<0.001, ****p<0.0001 GSIS for basal vs. glucose-stimulated condition using paired t-test. ^#^p<0.05, ^##^p<0.01, saline vs. Ex-4 treated mice using unpaired t-test. G) Percentage increase in insulin level compared to basal (prior to glucose gavage) insulin level. Mice treated chronically with Ex-4 show significantly reduced sensitivity to glucose for insulin secretion compared to saline treated mice (Week 9, ^#^p<0.05, GSIS in saline vs. Ex-4 treated mice using unpaired t-test).

### β-GASP1-KO do not develop tolerance to exendin-4

To determine whether GASP1 is critical for the development of tolerance to Ex-4 in intact mice, we repeated the same longitudinal paradigm as in Fig. 3 in β-GASP1-KO mice (Fig. 4A-F, Supp. Fig. 4). These data, summarized in Fig. 4G, show that β-GASP1-KO mice treated with Ex-4 do not develop tolerance to either exogenous Ex-4 (Fig. 4G, week 3 vs. week 8) or endogenous incretin (Fig. 4G, week 1 vs. week 9). When comparing the responses of WT vs. β-GASP1-KO mice, we found that the acute response of WT and β-GASP1-KO mice to both oral glucose (Fig. 4I, week 1) and Ex-4 (Fig. 4H, I week 2) are indistinguishable, while chronic treatment with Ex-4 produces substantial tolerance to Ex-4-stimulated insulin secretion in WT but not β-GASP1-KO mice (Fig. 4H, week 2 vs. 8). In addition, β-GASP1-KO treated chronically with Ex-4 show significantly greater GSIS compared to WT mice even in the absence of exogenous incretin drug (Fig. 4I, week 9).

**Figure 4:**
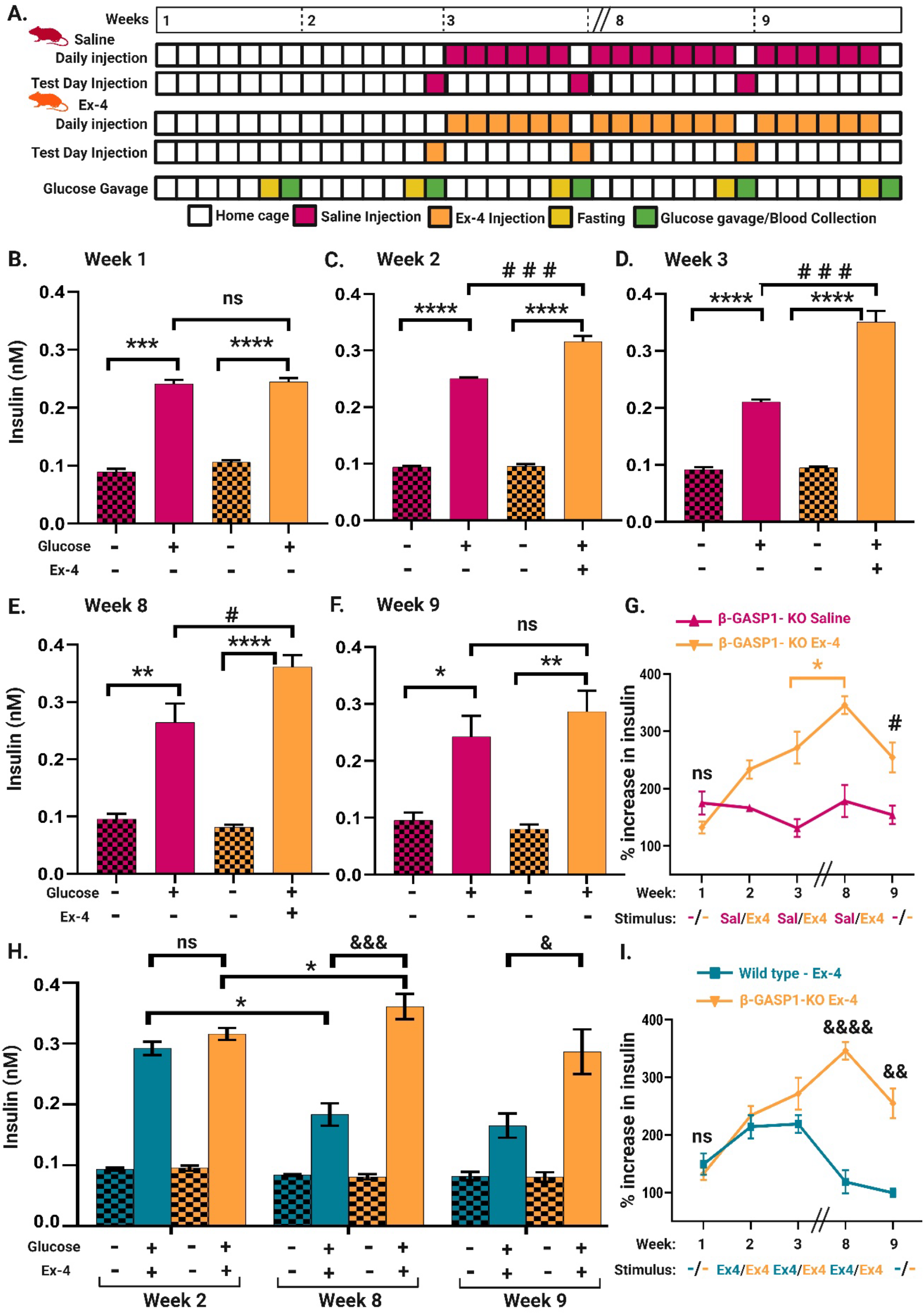
β-GASP1-KO mice do not develop tolerance to incretin. A) Schematic of the longitudinal mouse paradigm for monitoring plasma insulin levels in β-GASP1-KO mice treated with saline or Ex-4 (200μg/Kg). B) Glucose stimulated insulin secretion (GSIS) in β-GASP1-KO mice (n=9) at baseline (Week 1) following oral glucose administration (2g/kg body weight). β-GASP1-KO mice show robust GSIS in response to glucose gavage (hatched vs non-hatched bar, **p<0.001). C) GSIS in β-GASP1-KO mice following acute Ex-4 treatment (Week 2). Mice received either Ex-4 (n=5) or saline (n=4) injection on test day, followed by oral glucose administration and plasma insulin measurement. β-GASP1-KO mice treated with Ex-4 show significantly increased insulin secretion compared to saline treated mice (purple vs orange bar, ^###^p<0.001). D) GSIS in β-GASP1-KO mice following one week of either Ex-4 or saline treatment (Week 3). Mice were injected either with Ex-4 or saline twice daily for one week. Ex-4 treated mice maintain their Ex-4dependent incretin effect compared to saline treated mice (purple vs. orange bar, ^###^p<0.001). E). GSIS in β-GASP1-KO mice following six weeks of Ex-4 or saline treatment (Week 8). Mice continued to receive twice daily injection of either Ex-4 or saline for six weeks. β-GASP1-KO mice treated with Ex-4 maintain their incretin effect compared to saline treated mice (purple vs. orange bar, ^#^p<0.05) (no tolerance. F) GSIS in β-GASP1-KO mice following seven weeks of Ex-4 or saline treatment (Week 9). Mice continued to receive twice daily injections of either Ex-4 or saline. On test day, plasma insulin concentration was measured after oral glucose administration without either Ex-4 or saline injections. GSIS in mice chronically treated Ex-4 is comparable to saline treated mice (purple vs orange bar). The bar graphs (B-F) represent the mean ± S.E.M from multiple mice. *p<0.05, **p<0.01, ***p<0.001, ****p<0.0001 GSIS for basal vs. glucose-stimulated condition using paired t-test. ^#^p<0.05, ^###^p<0.001, saline vs. Ex-4 treated mice using unpaired t-test. G) Percentage increase in insulin level compared to basal level in saline vs. Ex-4 treated β-GASP1-KO mice. β-GASP1-KO mice do not show tolerance to Ex-4 after chronic Ex-4 treatment (*p<0.05, β-GASP1-KO Ex-4 mice week 3 vs 8, ^#^p<0.05, β-GASP1-KO saline vs Ex-4). H) GSIS comparing Ex-4 treated WT vs. β-GASP1-KO mice for week 2 – acute, week 8 – six weeks chronic and week 9 – seven weeks chronic Ex-4 treatment. Both WT and β-GASP1-KO mice show equivalent acute incretin effect (left panel). After six weeks of Ex-4 treatment, WT but not β-GASP1-KO mice show tolerance to Ex-4 (middle panel). WT mice show lower sensitivity to glucose for insulin secretion compared to β-GASP1-KO mice (right panel) after seven weeks of chronic Ex-4 treatment. The bar graphs (H) represent the mean ± S.E.M from multiple mice. ^&^p<0.05, ^&&^p<0.01, ^&&&^p<0.001, ^&&&&^p<0.0001 GSIS – Ex-4 treated WT vs. β-GASP1-KO using unpaired t-test. I). Percentage increase in insulin level compared to basal level in Ex-4 treated WT vs. β-GASP1-KO mice. β-GASP1-KO mice show no tolerance for Ex-4 dependent insulin secretion after chronic Ex-4 treatment and have high sensitivity to glucose for insulin secretion compared to WT mice. (Week 8 and 9, ^&&^p<0.01, ^&&&&^p<0.0001).

## Discussion

GLP-1R agonists have revolutionized the treatment of type 2 diabetes mellitus, offering improved glycemic control and other beneficial effects including weight loss and reduced cardiovascular risk^4,7,26^. Prolonged use can lead to reduced therapeutic response or tolerance to these important medications^22–25^, although the underlying mechanism for this diminishing therapeutic response is unknown. Here we show that tolerance to the incretin drug Ex-4 is dependent on GASP1, which mediates the post-endocytic sorting of GPCRs to the lysosome^38–40^. GASP1 disruption in either INS-1 cells or pancreatic beta cells did not affect acute GLP-1R signaling or insulin secretion *in vitro* or *ex vivo*. We also show that, with prolonged exposure to Ex-4, WT INS-1 cells (Fig. 1) and islets from WT mice (Fig. 2) showed diminished insulin secretion, while GASP1 knock-out INS-1 cells and islets maintained robust incretin-dependent insulin responses. WT mice likewise developed tolerance to Ex-4 during chronic treatment (Fig. 3), while mice with a selective disruption of GASP1 in pancreatic beta cells did not (Fig. 4). These findings underscore the crucial role of GASP1-mediated GLP-1R trafficking in the development of tolerance to incretin drugs and could suggest novel ways to improve therapeutic utility.

One way to prevent post-endocytic receptor degradation is to prevent receptor endocytosis^43^. GLP-1, Ex-4 and liraglutide all promote both G protein activation from the GLP-1R and arrestin recruitment and therefore GLP-1R endocytosis^30,44,45^. However, for many GPCR targets there are ligands that are differentially potent and/or efficacious at activating the G protein versus the arrestin signaling pathways. In particular, a ligand-specific signaling bias for the G protein versus the arrestin effector has been demonstrated for many classes of GPCR including the GLP-1R^46–48^. For example, tirzepatide (Mounjaro^®^) is a dual GLP-1R and glucose-dependent insulinotropic polypeptide (GIP) receptor agonist, which has been shown to have improved therapeutic utility, possibly through its action at both of these receptors^49–52^. Tirzepatide is also distinguished from other GLP-1R agonists in that it is biased for G protein signaling from the GLP-1R, as it doesn’t promote arrestin recruitment and receptor endocytosis^53,54^. By selectively biasing signaling towards insulin release and not arrestin recruitment, tirzepatide may allow the GLP-1R to evade GASP1-mediated post-endocytic degradation and thereby better sustain the efficacy of treatment. Since tirzepatide is not only biased for G protein at the GLP-1R but also strongly activates GIP receptors^52^, it is not possible to determine whether bias or dual agonism is key to its improved therapeutic effects. However, tirzepitide provides proof-of-concept that G-biased ligands at the GLP-1R are an achievable goal. In fact, several such molecules have been reported and shown to produce improved glycemic control in mice^44^.

GLP-1R agonist drugs have been approved more recently for weight management in individuals even without type-2 diabetes due to their ability to effectively regulate appetite, delay gastric emptying and impact central appetite control^55–57^. The long-term safety profile of GLP-1R agonists for weight loss in non-diabetic subjects is still being evaluated, as the use of these medications in the sole context of weight loss is relatively recent. Here, we observed that prolonged administration of Ex-4 to healthy non-diabetic WT mice resulted in a significant decrease in their responsiveness to glucose-stimulated insulin secretion during an oral glucose challenge when they were not on drug (Fig. 3G), indicating an altered response to their endogenous incretin hormones. This effect was not observed in the GASP1 knock-out mice (Fig. 4G, I). These findings suggest that caution should be exercised when considering the prolonged use of GLP-1R agonists as weight loss medications for healthy individuals, as prolonged treatment with these agonists has the potential to modify the body’s insulin secretion profile and disrupt glucose homeostasis.

The present study highlights the crucial role of GASP1 in regulating GLP-1R responses during chronic drug administration that mimic their intended therapeutic use. Our observations suggest that novel ligands with a bias for G protein over arrestin signaling could have longer therapeutic efficacy in type 2 diabetes patients. G biased ligands might also be safer for use long term for weight loss as they would not produce tolerance and subsequent loss of insulin secretion from pancreatic islets that we attribute here to GASP1-dependent signaling.

## Methods

### Cell Culture

Rat insulinoma-derived insulin producing INS-1 cells were purchased from AddexBio (C0018007). Cells were cultured using AddexBio optimized RPMI -1640 media (C0004-02) containing 11mM glucose and L-glutamine, supplemented with 10% fetal bovine serum (Corning, 35-010-CV) and 0.05 mM 2-mercaptoethanol (Sigma-Aldrich, M6250). INS-1 cells were sub-cultured using 0.25% trypsin/EDTA (Corning, 25-053-CI) when 70-80% confluency was reached. HEK293 and HEK293T cells were purchased from ATCC and cultured in DMEM media (Gibco, 11965-092) containing L-glutamine and 4.5 g/L D-glucose, supplemented with 10% FBS. Both INS-1 and HEK293 cells were cultured under standard aseptic tissue culture conditions and maintained at 37°C in 5% CO_2_ humidifier incubator.

### CRISPR/Cas9 mediated gene targeting

Single guide RNAs (sgRNA) targeting the rat GPRASP1 (GASP1 protein) gene were purchased from Applied Biological Materials (ABM, 22601116). The sgRNAs target three distinct sites of the GPRASP1 gene and were cloned into a lentiviral vector system (pLenti-U6). Lentiviruses expressing Cas9 and sgRNAs were produced by co-transfecting HEK293T cells with sgRNAs and packaging plasmids (ABM, LV003) as per the manufacturer’s protocol. Supernatant containing lentivirus was harvested after 48-hours. The viral supernatant was filtered using a 0.45-micron filter (Nalgene, 190-2545) to remove any HEK293T cells and concentrated by centrifuging at 25,000 RPM for 100 minutes at 37°C. The viral concentrate was resuspended in complete RPMI – 1640 media and used for transduction of INS-1 cells. Hexadimethrine bromide (Polybrene, sc134220) was also added to the cells at 2 μg/mL concentration to enhance lentiviral infection efficiency. 48-hours post-transduction, puromycin (Invitrogen, A11138) was added at 2 μg/mL concentration for 7-10 days. After selection in puromycin, cells were allowed to grow until visible colonies formed. Colonies derived from single cells were picked, expanded and examined for GASP1 protein expression.

### Validation of CRISPR/Cas9 deletion proficiency

Disruption of the GPRASP1 gene was evaluated using immunoblot analysis for the GASP1 protein. Cells from single cell-derived colonies were lysed in RIPA lysis buffer (Thermofisher Scientific, 89900) containing an EDTA-free protease inhibitor mini tablet (Roche, 11836170001). The lysates were resolved by SDS-PAGE and transferred to PVDF membrane (Bio-Rad, 1620177). The membrane was probed for GASP1 expression with a rabbit polyclonal anti-GASP1/2 antibody we generated (1/1000)^58^. GAPDH was used as a loading control (Invitrogen, MA5-15798). Immune complexes were detected using IRDye® conjugated secondary antibodies-goat anti-rabbit 800CW (Odyssey, 926-32211) and goat anti-mouse 680LT (Odyssey, 926-68020) respectively. The images were developed using the Odyssey CLx Li-Cor system.

### Intracellular cAMP homogenous time-resolved fluorescence (HTFR) Assay

INS-1 cells were seeded in a 384-well white, low volume, flat bottom plate (Fisher scientific, 781981) at 7 × 10^3^ cells/well and incubated overnight in AddexBio optimized RPMI-1640 media. Cells were washed twice with PBS and pretreated either with vehicle (media) or 100nM Ex-4 (Tocris, 1933) for 3-hours. Following pretreatment, cells were washed and allowed to recover for 15 minutes and then stimulated with either vehicle (PBS) or increasing concentrations of Ex-4 diluted in PBS containing 100 μM of IBMX (Sigma-Aldrich, I5879) for 30 minutes at 37°C. Cells were lysed and intracellular cAMP was measured using an HTRF immunoassay (CisBio - cAMP G_s_ dynamic kit, 62AM4PEB) according to the manufacturer’s instructions and measured using a Flexstation-3 (Molecular Devices). The concentration of cAMP (nM) in each sample was extrapolated from a cAMP standard curve. The results were expressed as cAMP dose-response curves fitted using non-linear regression in GraphPad Prism 9. For determination of E_max_ in Supp. Table 1, WT-vehicle pretreated cells treated with 100nM of Ex-4 was defined as 100% and all other samples normalized to this treatment.

### Insulin secretion assay in INS-1 cells

INS-1 cells were seeded in a 384-well white, low volume, flat bottom plate (Fisher Scientific, 781981) at 7 × 10^3^ cells/well and incubated overnight in AddexBio optimized RPMI-1640 media. Cells were washed twice with Krebs-Ringer Bicarbonate buffer (KRB buffer, 130mM NaCl, 5mM KCl, 1.2mM CaCl_2_, 1.2mM MgCl_2_, 1.2mM KH_2_PO_4_, 25mM NaHCO_3_, 20mM Hepes pH7.4) and glucose starved with RPMI media containing 10% FBS and 1% penicillin/streptomycin for 3-hours. During starvation, cells were either treated with vehicle (media) or 100nM Ex-4 (Tocris, 1933). After glucose starvation and pretreatment, cells were washed with KRB buffer and recovered in RPMI media containing 5.5mM glucose for 15 minutes. Following recovery, cells were stimulated with 11mM glucose ± increasing concentrations of Ex-4 in KRB buffer containing 0.1% w/v BSA for 30 minutes. After treatment, supernatant containing secreted insulin was collected and transferred into a new 384-well plate. The insulin concentration was determined using an insulin HTRF assay kit (CisBio, 62IN1PEG) according to manufacturer’s instructions and measured using a Flexstation-3 (Molecular Devices). The amount of insulin (ng/mL) in each sample was extrapolated from an insulin standard curve. The results were expressed as insulin dose-response curves fitted using non-linear regression in GraphPad Prism 9. For the E_max_ calculations in Supp. Table 2, WT-vehicle pretreated cells treated with 100nM of Ex-4 was defined as 100% and all other samples normalized to this treatment.

### Mice

Mice were bred in-house and handled according to National Institute of Health guidelines for the care and use of laboratory animals. All the protocols were approved by the Institutional Animal Care and Use Committee (IACUC) at the University of California, Davis. Adequate measures were taken to minimize animal suffering and discomfort. Mice were housed 2-5 per cage in temperature- and humidity-controlled rooms with 12:12 hours light:dark sleep cycle and provided with food and water *ad libidum*.

### Generation of Floxed-GASP1 conditional knockout mice

Supplemental Fig. 1A is a schematic of the GPRASP1 locus on the mouse X chromosome. A targeting vector was designed with a neomycin (G418)-resistance gene flanked by loxP sites inserted into intron 4 upstream of the GPRASP1 gene and a third loxP site inserted downstream of the GASP1 open reading frame (Supp. Fig. 1B). The linearized targeting vector was electroporated into ∼10^7^ C57BL/6 ES cells and clones were selected with 200μg/ml G418. ES cells with homologous recombination of the targeting vector (Supp. Fig. 1C) were determined by Southern blot. These were treated with Cre recombinase and clones where loxP sites 1 and 2 were recombined were identified by PCR (Supp. Fig. 1D). Clones were implanted into C57/Bl6 mothers and germline transmission of the fl-GASP1 conditional KO gene was confirmed by breeding. Genotyping is performed with a set of 3 primers (5’ – 3’ sequence):

WT forward primer:

GAGTGACTACTGTGAGACTTGG

GASP1-KO forward primer:

GTGAACTGAGCCGTTGTAAATAAGATGC

Common reverse primer:

CATCTCTTCGATTTATAGTTCTCCCACC

### Generation of β-cell specific GASP1 knock out mice

To generate pancreatic β-cell specific GASP1 knock-out mice (β-GASP1-KO) we bred floxed GASP1 mice (Supp. Fig. 1)^59^ to *UCN3-Cre* driver mice^42^. GASP1 is an X-linked gene. Female homozygous floxed GASP1 mice (GASP1^fl/fl^) were crossed with male hemizygous *UCN3-Cre* driver mice (*UCN3*^Cre/+^). Male littermate from this cross were genotyped for the floxed GASP1 allele and the *UCN3-Cre* transgene using tail DNA. Mice with and without the *UCN3-Cre* transgene were used in all experiments (Fig. 2A). The following primers were used for genotyping of these mice (5’ – 3’ sequence):

GASP1:

WT forward primer:

GAGTGACTACTGTGAGACTTGG

GASP1-KO forward primer:

GTGAACTGAGCCGTTGTAAATAAGATGC

Common reverse primer:

CATCTCTTCGATTTATAGTTCTCCCACC

UCN3:

Forward: CGAAGTCCCTCTCACACCTGGTT

Reverse: CGGCAAACGGACAGAAGCATT

### Pancreatic Islet isolation

Mouse pancreatic islets were obtained from 10-12 weeks old C57BL/6 WT or β-GASP1-KO mice. Islets were isolated by infusing the pancreatic duct with HBSS (no Ca^2+^ or Mg^2+^, Gibco, 14170-112) containing 0.8 mg/mL collagenase P (Roche, 11249002001). The pancreata were dissected and digested at 37°C for 13 minutes in a water bath. Islets were subsequently isolated using a Histopaque gradient (Sigma, 10771). The isolated islets were further purified by picking twice into fresh HBSS + 5% NCS (Gibco, 16010-167) + 1mM CaCl_2_ (Sigma-Aldrich, C5080). Isolated islets recovered overnight in RPMI-1640 supplemented with 10% FBS and 1% penicillin/streptomycin (Gibco, 15140-122)^41^.

### mRNA extraction and cDNA synthesis

Total RNA was extracted from isolated WT or β-GASP1-KO islets using a phenol:chloroform:isopropanol extraction method. Briefly, islets were collected into an RNase-free microcentrifuge tube containing 500μL Trizol reagent (Invitrogen,15596018) followed by addition of 100μL of chloroform (Fisher Scientific, C298-500). The tubes were shaken for 20 seconds and centrifuged at 12,000g for 15 minutes at 4°C. The clear top aqueous phase containing RNA was collected into a new RNase-free tube. RNA was precipitated by adding 250μL of isopropanol (Fisher Scientific, BP2618500) and centrifuging the tube at 12,000g for 10 minutes at 4°C. The RNA pellet was then washed with ethanol, air dried and re-suspended in RNase free water. The cDNA was synthesized from extracted RNA by reverse transcription using high-capacity RNA-to-cDNA™ kit (Applied Biosystems, 4368813) according to manufacturer’s instructions.

### Quantitative real-time PCR

cDNA synthesized from RNA extracted from WT or β-GASP1-KO islets were used to determine GPRASP1 and GLP-1R expression in WT or β-GASP1-KO islets using quantitative real-time PCR (qPCR). The cDNA was amplified using PowerUp™ SYBR™ green super mix (Applied Biosystems, A25742) through 30-40 cycles of qPCR. For the qPCR reaction we used 0.5μM forward and reverse primer. For GPRASP1, forward primer: 5’-TGGTTCTGGGCAGATGATGAAGAGA-3’ and reverse primer: 5’-TTGTTGCTTTTGTAGATGCCGACC-3’ were used. For GLP-1R, forward primer: 5’-CCCTGGGCCAGTAGTGTG-3’ and reverse primer: 5’-GCAGGCTGGAGTTGTCCTTA-3’ were used. Quantification was performed using the 2^(ΔΔCt) method. Data were normalized to an endogenous housekeeping gene (hypoxanthine phosphoribosyltransferase 1, HPRT gene) and expressed as relative gene expressions compare to WT control islets. For the HPRT gene, forward primer: 5’-TCCTCCTCAGACCGCTTTT-3’ and reverse primer: 5’-GCAGGCTGGAGTTGTCCTTA-3’ were used.

### Mouse islet glucose stimulated insulin secretion (GSIS) assay

Following overnight recovery after isolation, mouse islets were picked twice in KRB buffer (130mM NaCl, 5mM KCl, 1.2mM CaCl_2_, 1.2mM MgCl_2_, 1.2mM KH_2_PO_4_, 25mM NaHCO_3_, 20mM HEPES pH7.4) supplemented with 0.1% BSA and 5.5mM glucose. The islets were placed in a 48-well plate and glucose starved for 1-hour at 37°C in KRB + 0.1% BSA + 5.5mM glucose. The islets were then stimulated for 1-hour at 37°C with KRB buffer supplemented with low glucose (5.5mM), high glucose (11mM) or high glucose + 100nM Ex-4 (Tocris, 1933). After an hour, supernatants were collected for insulin detection (Acute, Fig. 2C). The islets were then treated with 100nM Ex-4 diluted in complete RPMI-1640 media for 3-hours at 37°C. Following Ex-4 treatment, the islets were washed with KRB buffer and allowed to recover for 15 minutes. They were then treated with KRB supplemented with low glucose, high glucose and high glucose + Ex-4 for 1-hour and supernatants were collected for insulin detection (Tolerance, Fig 2C). The islets were then washed with KRB and placed in RPMI – 1640 complete media and allowed to recover. After 24-hours, the islets were again glucose starved for an hour followed by stimulation with KRB buffer supplemented with low glucose, high glucose with or without Ex-4 for 1-hour. The supernatants were collected after incubation for insulin detection (Recovery, Fig. 2C).

### Oral glucose tolerance test (OGTT) and blood collection

Blood was collected for OGTT and plasma insulin measurements from age-matched WT and β-GASP1-KO mice. A total of 8 WT and 9 β-GASP1-KO mice were randomly assigned into two experimental groups - control (saline) and treatment (Ex-4): WT saline (n=4), WT Ex-4 (n=4), β-GASP1-KO saline (n=4) and β-GASP1- KO Ex-4 (n=5). The mice were bred in-house and handled gently to minimize stress. Mice were entered into the longitudinal study paradigm (Fig. 3A and 4A, Supp. Figs. 3 and 4) at 8 weeks old. At week 1, baseline glucose tolerance and plasma insulin were measured. Mice were fasted overnight (∼14-16 hours) and then administered a 2g/kg oral bolus of glucose. Blood glucose was measured from blood using standard glucometer (TrueFocus, Walgreen) at 0.5-,15-,30-,60-,90- and 120 minutes post glucose gavage. Blood (∼100μL) was also collected from the tail vein of each mouse into a microvette capillary tube (Thermofisher, 16.444.100) at 0 and 15 minutes after glucose administration and centrifuged at 1500xg for 10 minutes at 4°C to prepare plasma (Supp. Fig. 2A). The plasma samples were stored at -80°C until analysis. The mice were returned to their home cages for a week where they had access to water and food *ad-libidum*. At week 2, after overnight fasting, mice were injected either with saline or 200μg/Kg Ex-4 (MedChemExpress,HY-13443) 15 minutes prior to OGTT. After 15 minutes, mice received an oral glucose gavage at the 0 time-point. Blood glucose levels were measured at 5-,15-,30-,60-,90- and 120 minutes after glucose administration (Supp. Fig. 3B). Blood samples (∼100μL) were collected for plasma insulin measurement before the saline or Ex-4 injection and 15 minutes after glucose gavage (Acute Ex-4 treatment) (Supp. Fig. 2B). The mice were returned to their home cage and injected twice daily with either saline or Ex-4 (200μg/Kg) for an an additional seven weeks. Oral glucose tolerance test and blood plasma collection were performed at week 3 and 8 (1 and 6 weeks of chronic Ex-4 treatment respectively) (Supplemental figure S2B). On week 9 (7 weeks of chronic Ex-4 treatment), the mice were fasted overnight and administered a 2g/Kg oral bolus of glucose without prior saline or Ex-4 injection. Oral glucose tolerance was measured over a period of 120 minutes and blood was collected for plasma insulin at 0- and 15 minutes post oral gavage (Supp. Fig. 2A).

### Insulin measurement from GSIS islet study and mouse plasma

The insulin secreted from islets ex vivo and plasma insulin from blood samples a were measured using Lumit Insulin Immunoassay kit (Promega, CS3037A05) as per the manufacturer’s instructions and measured using a Flexstation-3.

### Statistics

Data are presented as means ± SEM and the number of experiments is indicated in each experiment. GraphPad Prism version 9 was used to perform all statistical analysis. The student’s t-test or two-way analysis of variance (ANOVA) with Tukey’s multiple comparison test was performed in GraphPad Prism to detect statistical differences. P < 0.05 was considered statistically significant. For the E_max_ and EC_50_ determination, vehicle pretreated WT-cells treated with 100nM Ex-4 was set at a reference value of 100%. All other samples were normalized to this treatment. The normalized data is then used to determine the E_max_ and EC_50_ values using GraphPad Prism version 9.

## Acknowledgements

The author would like to thank all members of the J. Whistler lab for their valuable input and ongoing support for this research. The author would also like to thank members of Huising lab for their support. This study was funded by American Diabetes Association – Precision Medicine & Diabetes Innovative Basic Science Award (Award number: 7-22-IBSPM-06), Stanford Diabetes Research Center Grant and by the funds provided by the state of California as a start up to J.L.W.

## Authors Contributions

J.L.W and A.G design, conceptualize and investigate the experiments. A.G optimized the experimental protocols. A.G, M.K, N.D, and M.F performed the experiments. Data analysis was performed by A.G, C.M.K and J.L.W. J.L.W and A.G designed and generated the figures. J.L.W supervised and administered the project as well as provided the resources for the project. B.P.C and M.O.H provide necessary training and supervision required for *ex vivo* and *in vivo* experiments. A.G and J.L.W wrote the manuscript. All authors contributed to the editing of the manuscript.

## Declaration of Interest

The authors declare no competing interests.

## Inclusion and Diversity

We support inclusive, diverse, and equitable conduct of research.

## Supplemental Figures and Tables

**Supplemental Figure 1:**
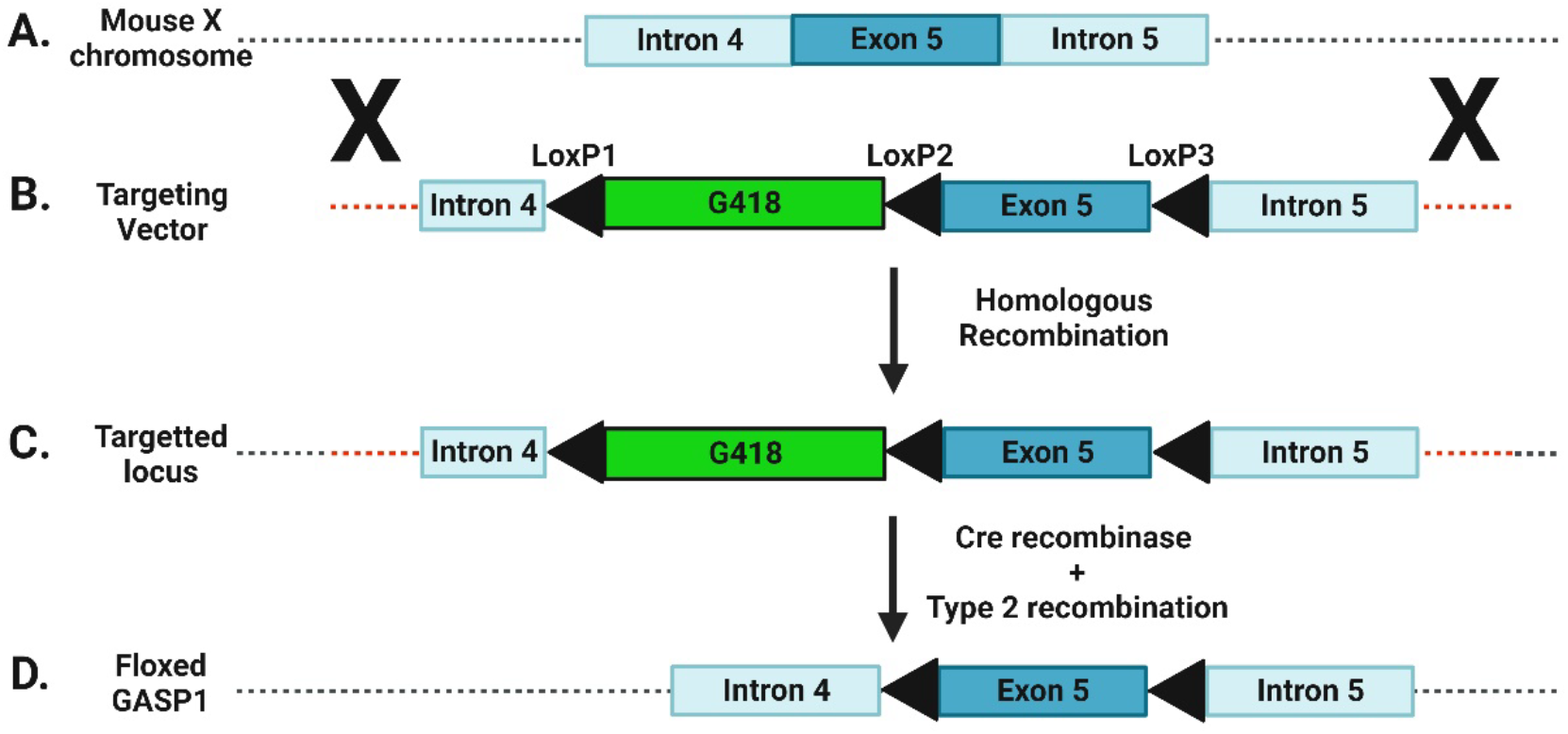
Generation of flox-GASP1 mice. A) The GASP1 gene is located on the mouse X chromosome. The open reading frame (ORF) for the GASP1 protein is encoded by a single exon (Exon 5). B) A targeting vector was constructed from this locus in which G418 flanked by lox P sites (lox P site 1 and 2) was inserted in the intron upstream of the GASP1 ORF, and a third lox P site (lox P site 3) was inserted downstream of the GASP1 ORF. C) The targeting vector was electroporated into ES cells of C57/Bl6 mice and clones selected by neomycin resistance. Homologous recombination into the GASP1 locus was detected from individual clones by Southern hybridization. D) Individual clones were incubated with Cre recombinase and clones where only loxP sites 1 and 2 were recombined were identified by PCR screening to create the floxed GASP1 locus.

**Supplemental Figure 2:**
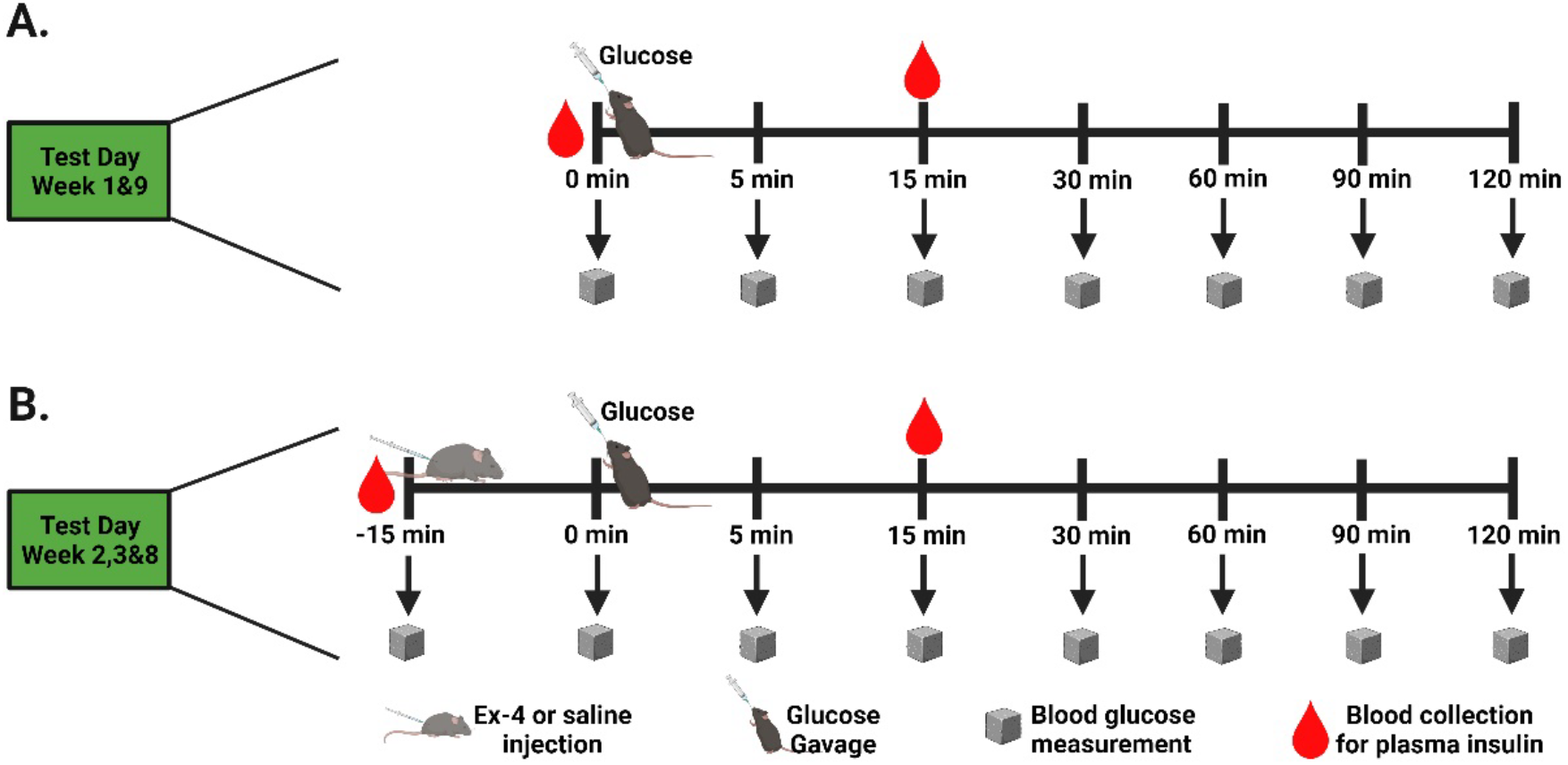
Schematic illustration of sample collection during the oral glucose tolerance test and plasma insulin measurement in mice. A) Schematic of the oral glucose tolerance test (OGTT) performed in mice on week 1 & 9. Mice received oral glucose (2g/Kg body weight) at the 0 time-point through glucose gavage. Blood glucose levels were measured at 5-, 15-, 30-, 60-, 90- and 120- minutes after oral glucose administration. Blood samples were also collected for plasma insulin measurement before glucose administration and 15 minutes post glucose gavage. B). Schematic of the OGTT performed in mice on week 2,3 & 8. Mice were injected either with saline or Ex-4 (200 μg/Kg) 15 minutes prior to glucose gavage. Mice received oral glucose at 0 time-point. Blood glucose levels were measured for 120 minutes. Blood samples were collected for plasma insulin measurement before saline or Ex-4 injection and 15 minutes post glucose gavage.

**Supplemental Figure 3:**
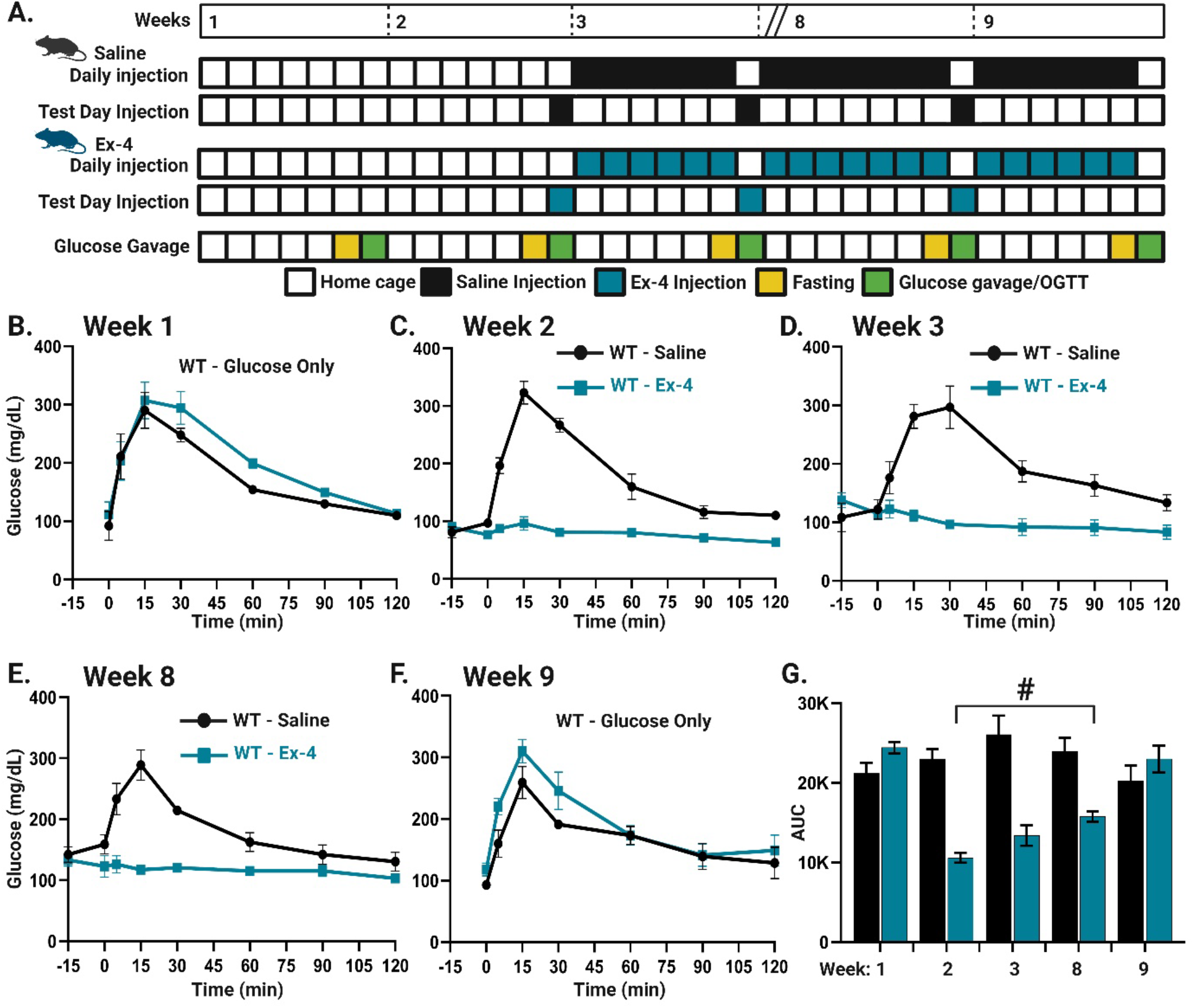
Glucose clearance in wild-type mice. A) Schematic of the longitudinal mouse paradigm for the OGTT in WT mice treated with saline or Ex-4 (200μg/Kg). B) OGTT in WT mice (n=8) at baseline (Week 1). Blood glucose levels were measured at the time-points shown after oral glucose administration (2g/kg body weight). C) OGTT in WT mice after acute Ex-4 treatment (Week 2). Blood glucose levels were measured after oral glucose challenge in WT mice following saline (n=4) and Ex-4 (n=4) treatment. Ex-4 treated mice show significantly faster clearance of glucose compared to saline treated mice (****p<0.0001). D) OGTT in WT mice following one week of twice daily Ex-4 or saline treatment (Week 3). Ex-4 treated mice show significantly faster clearance of glucose compared to saline treated mice (****p<0.0001). E) OGTT in WT mice after six weeks of chronic Ex-4 or saline treatment (Week 8). Mice continue to receive twice daily injections of either Ex-4 or saline for six weeks. Ex-4 treated mice continue to show significantly faster clearance of glucose compared to saline treated mice (*p<0.05), however, WT Ex-4 treated mice exhibit significantly lower AUC values at week 2 vs week 8 (^#^p<0.05, Panel G). F) OGTT in WT mice after seven weeks of chronic Ex-4 or saline treatment (Week 9). Mice continue to receive twice daily injections of either Ex-4 or saline. On test day, blood glucose levels were measured at the time points shown after oral glucose administration without saline or Ex-4 injection. Representative OGTT curves (B-F) showing mean ± standard error of mean glucose level. G) Bar graph representing the calculated area under the curves values (AUC) in WT mice treated with or without Ex-4 (n=4 mice per group). Statistical significance was determined using two-way ANOVA followed by post-hoc Tukey’s multiple comparison test (*p<0.05, ****p<0.0001 Saline vs Ex-4 treatment, ^#^p<0.05 Ex-4 treated mice week 2 vs week 8).

**Supplemental Figure 4:**
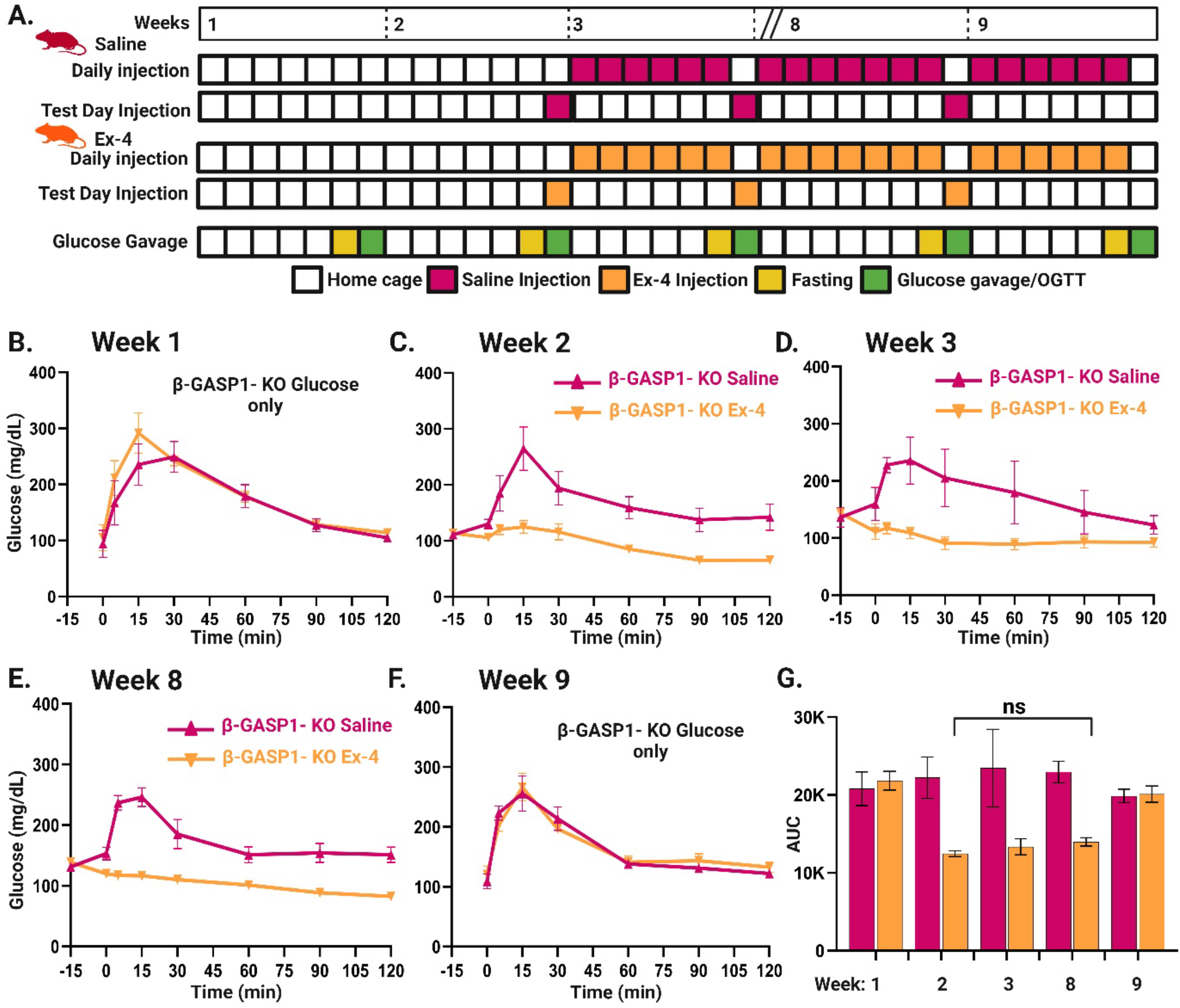
Glucose clearance in β-GASP1-KO mice. A) Schematic of the longitudinal mouse paradigm for the OGTT in β-GASP1-KO treated with saline or Ex-4 (200μg/Kg). B) OGTT in β-GASP1-KO mice (n=9) at baseline (Week 1). Blood glucose levels were measured at the time points shown after oral glucose administration (2g/kg body weight). C) OGTT in β-GASP1-KO mice after acute Ex-4 treatment (Week 2). Blood glucose levels were measured after oral glucose challenge in β-GASP1-KO mice following saline (n=4) and Ex-4 (n=5) treatment. Ex-4 treated mice show significantly faster clearance of glucose compared to saline treated mice (*p<0.05). D) OGTT in β-GASP1-KO mice following one week of twice daily Ex-4 or saline treatment (Week 3). Ex-4 treated mice show significantly faster clearance of glucose compared to saline treated mice (*p<0.05). E) OGTT in β-GASP1-KO mice after six weeks of chronic Ex-4 or saline treatment (Week 8). Mice continue to receive twice daily injections of either Ex-4 or saline for six weeks. Ex-4 treated mice show significantly faster clearance of glucose compared to saline treated mice (*p<0.05). There is no tolerance to Ex-4 in β-GASP1-KO mice after six weeks chronic Ex-4 treatment (Panel G, Ex-4 treated β-GASP1-KO week 2 vs week 8). F) OGTT in β-GASP1-KO mice after seven weeks of chronic Ex-4 or saline treatment (Week 9). Mice continued to receive twice daily injections of either Ex-4 or saline. On test day, blood glucose levels were measured at time points show after oral glucose administration without saline or Ex-4 injection. Representative OGTT curves (B-F) showing mean ± standard error of mean glucose level. G) Bar graph representing the calculated area under the curves values (AUC) in β-GASP1-KO mice treated with (n=5) or without (n=4) Ex-4. Statistical significance was determined using two-way ANOVA followed by post-hoc Tukey’s multiple comparison test (*p<0.05, Saline vs Ex-4 treatment).

**Table S1:** Effect of Ex-4 pretreatment on cAMP accumulation in GASP1 WT vs KO INS-1 cells.

| INS-1 cells | Pretreatment | E<sub>max</sub> (%) | EC<sub>50</sub> (nM) |
| --- | --- | --- | --- |
| GASP1 -WT | Vehicle | 100 | 0.59 ± 0.28 |
|  | Ex-4 | 44.30 ± 11.53 ** | 0.78 ± 0.42 |
| GASP1-KO | Vehicle | 95.14 ± 3.26 | 0.55 ± 0.26 |
|  | Ex-4 | 92.17 ± 4.32 <sup>ns</sup> | 0.76 ± 0.34 |

<sup>a</sup> The E<sub>max</sub> and EC<sub>50</sub> values were calculated using Figure 1 in Prism. The data is shown as mean ± S.E.M.
<sup>b</sup> Cyclic AMP accumulation was measured in rat INS-1 cells with or without GASP1.
<sup>c</sup> \*\* p < 0.01, ns = not significant comparing vehicle pretreatment vs Ex-4 pretreatment using unpaired t - test.

**Table S2:**
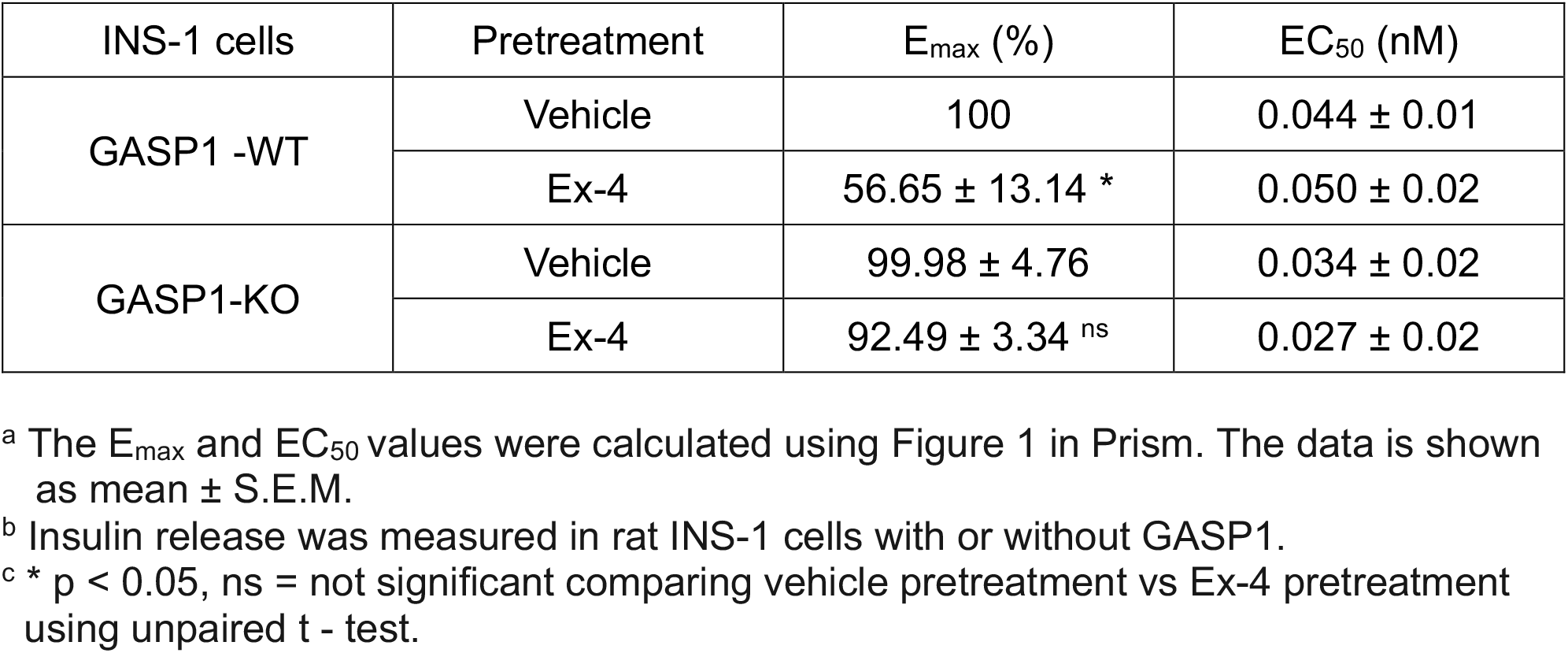
Effect of Ex-4 pretreatment on insulin release in GASP1 WT vs KO INS-1 cells.

## Notes

### Competing Interest Statement

The authors have declared no competing interest.

